# AntAngioCOOL: An R Package for Computational Detection of Anti-Angiogenic Peptides

**DOI:** 10.1101/233601

**Authors:** Javad Zahiri, Babak Khorsand-Ghaffari, Ramin Shirali Hossein Zade, Mohammadjavad Kargar, Ali Akbar Yousefi

## Abstract

Angiogenesis inhibition research is a cutting edge in angiogenesis-dependent disease therapy, and especially in cancer therapy. Recently, studies on anti-angiogenic peptides have provided promising results in the cancer treatment field. In the current study we propose an effective machine learning based R package (AntAngioCOOL) to predict anti-angiogenic peptides. We have examined more than 200 different classifiers to build an efficient predictor. Also, more than 17000 features have been extracted to encode the peptides. However, finally, more than 2000 informative features have been selected to train the classifiers. According to the obtained results AntAngioCOOL can effectively predict anti-angiogenic peptides: this tool achieved sensitivity of 88%, specificity of 77% and accuracy of 75% on independent test set. AntAngioCOOL can be accessed at https://cran.r-project.org/.

## INTRODUCTION

Angiogenesis is the process of formation of new blood vessels from pre-existing vessels to make a supply of nutrients and a waste disposal pathway1. Angiogenesis is a normal and fundamental physiological process in growth and development 2-4. However, it is a vital event in cancer progression- transition tumors from a benign state to a malignant one- and spread of a tumor (metastasis)5-7. Nowdays, decreasing or inhibiting angiogenesis is a cutting edge research in cancer therapy and is in the center of other angiogenesis-dependent disease therapy1,8-15.

Recognition the anti-angiogenic peptides has stimulated great interest among researchers in the cancer treatment field during recent years 16, as well as other therapeutic peptides 17-23. However, there exist very rare studies in computational detection of ant-angiogenic peptides8.

In this paper, we have proposed an efficient machine learning based R package to detect anti-angiogenic peptides, namely AntAngioCOOL. Five types of features have been used to encode peptides in order to predict anti-angiogenic ones. According to the obtained results, AntAngioCOOL reached to a satisfactory performance in anti-angiogenic peptide prediction on a benchmark non-redundant independent test dataset.

## METHODS

### Dataset

We have used the gold standard dataset that has been recently published ^8^. After removing redundant peptides, this dataset contains 135 anti-angiogenic peptides (positive instances) and 135 non anti-angiogenic peptides (negative instances). Also, a 20% of each class has been selected to construct an independent test dataset (see supplementary file).

### Features

The following subsections provide a brief description for each peptide feature. Moreover, table 1 demonstrates the distribution of the features that have been used to encode each peptide.

**Table 1:** Distribution of the features used to encode each peptide.

| Feature type |  | No. of features |
| --- | --- | --- |
| PseAAC($\lambda=6$) | | 28 |
| k-mer composition | k=2 | 400 |
|  | k=3 | 8000 |
|  | k=4 | 160000 |
| k-mer composition<br>(reduced alphabet*) | k=2 | 64 |
|  | k=3 | 512 |
|  | k=4 | 4096 |
| physico-chemical profile |  | 1910 |
| atomic profile |  | 80 |
| Total |  | 175062 |

\*To compute k-mer composition features, the reduced amino acid alphabet proposed by Zahiri et al has been applied: the 20
alphabet of amino acids have been reduced to a new alphabet with size 8 according to 544 physicochemical and biochemical indices that extracted from AAIndex database (C1={A, E}, C2={I, L, F, M, V}, C3={N, D, T, S}, C4={G}, C5={P}, C6={R, K, Q, H}, C7={Y, W}, C8={C}). We have computed k-mer composition for k=2,3,4 for each peptide.

### Pseudo Amino Acid Composition

We used pseudo amino acid composition (PseAAC) that has been used effectively in predicting cell penetrating peptides 21. Despite the simple amino acid composition, PseAAC considers the sequence-order information of the peptide. Interested readers may refer to 24 for more information on PseAAC.

### k-mer composition

k-mer composition shows the fraction of all possible subsequences with length k in the given peptide. Also, the reduced amino acid alphabet proposed by Zahiri et al 25 has been applied to compute another k-mer composition: the 20 alphabet of amino acids have been reduced to a new alphabet with size 8 according to 544 physicochemical and biochemical indices that extracted from AAIndex database 26 (C1={A, E}, C2={I, L, F, M, V}, C3={N, D, T, S}, C4={G}, C5={P}, C6={R, K, Q, H}, C7={Y, W}, C8={C}). We have computed k-mer compositions for k=2,3,4 for each peptide.

### Physico-chemical profile

In order To compute this feature type, 544 different physico-chemical indices have been extracted from AAIndex 26. To remove redundancies, a subset of indices with correlation coefficient less than 0.8 and greater than -0.8 has been selected, which resulted in 191 non-redundant physico-chemical indices.

This feature type has been extracted for 5 amino acids of N- termini (5-NT) and C-termini (5-CT). Finally, each peptide has been encoded as a 10×191-dimentional feature vector as below:

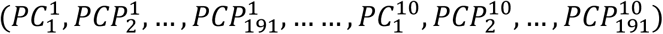

where 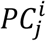 is the value of the jth physico-chemical index for the ith amino acid of the peptide (for i=1..5 in the 5-CT and i=6..10 in 5-NT)

### Atomic profile

A 50-dimentional feature vector has been used to encode each peptide according to its atomic properties as below:

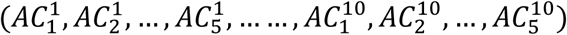

where 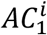 through 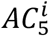 represent the frequency of five types of atoms: C, H, N, O, S in the ith amino acid of the peptides (for i=1..5 in the 5-CT and i=6..10 in 5-NT). For details of atomic composition for each 20 natural amino acid see 17.

### Machine Learning Method

To build a powerful anti-angiogenic peptide predictor, 227 different classifiers (see supplementary file) in the caret package 27 have been examined. Finally, the three best classifiers (those with best sensitivity, specificity and accuracy) have been selected to be included in the AntAngioCOOL package. Figure 1 provides a schematic representation of the proposed method.

**Figure 1:**
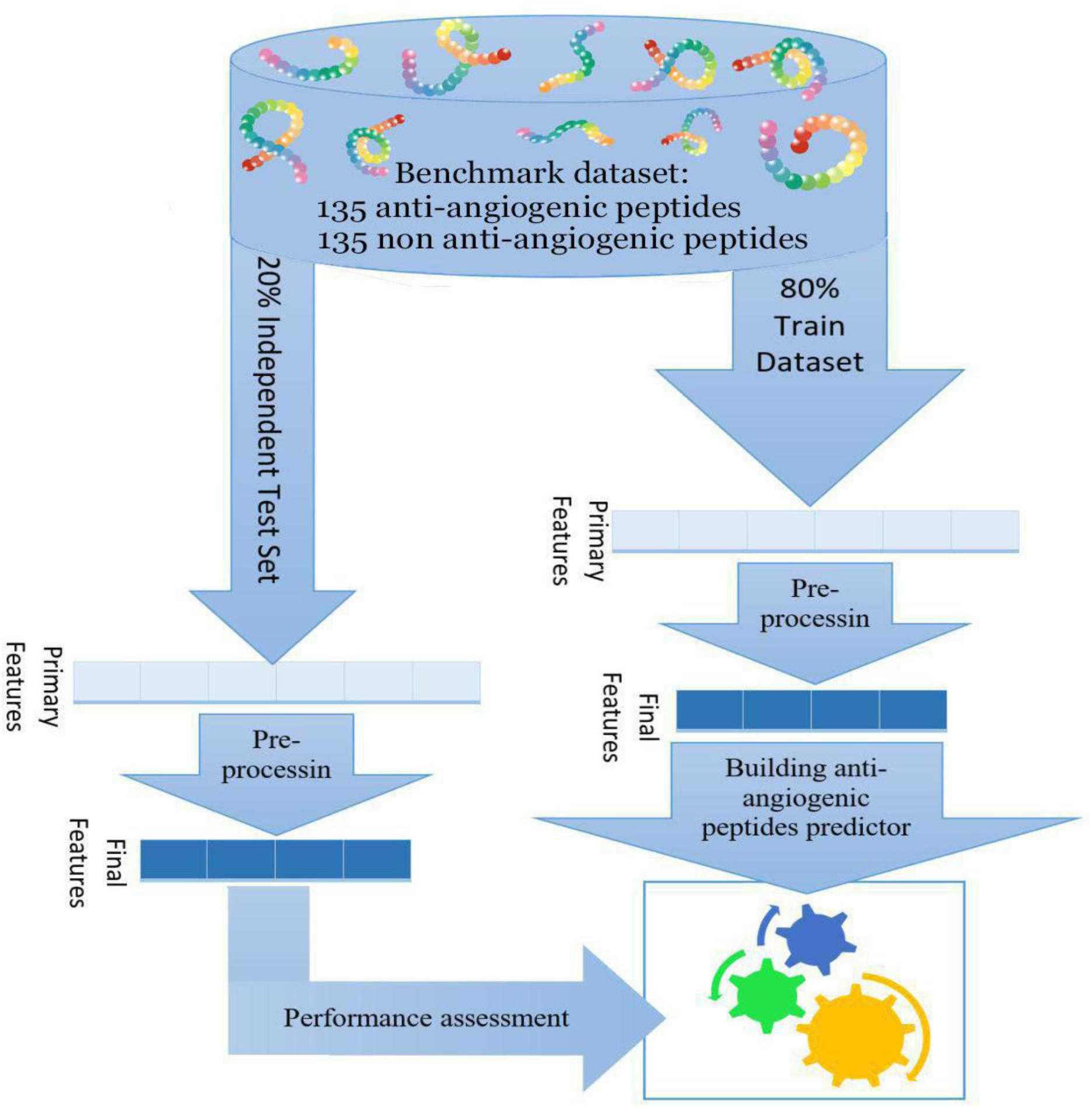
Schematic representation of the proposed method (AntAngioCOOL) for anti-angiogenic peptide prediction.

### Evaluation parameters for the prediction performance

The training dataset was used to train the classifier, and then the classifier was evaluated using the test data. The predictions made for the test instances were used to compute the following performance measures:

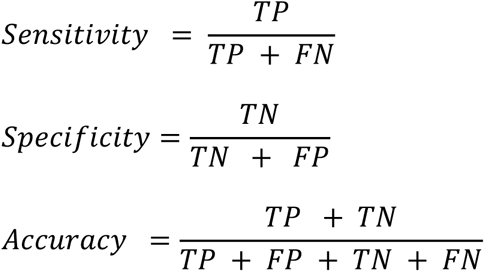

Where, TP and TN are the number of correctly predicted anti-angiogenic peptides and non anti-angiogenic peptides, respectively. Similarly, FP and FN are the number of non anti-angiogenic peptides and anti-angiogenic peptides that wrongly predicted as anti-angiogenic peptides and non anti-angiogenic peptides, respectively.

## 3. RESULTS & DISCUSSION

### 3.1 Preprocessing

To remove non-informative features, which can lead to reducing the computational cost without losing the prediction performance, nearZeroVar function from caret package27 has been utilized. This function eliminates those features that have one unique value (i.e. are zero variance features) or features with both of the following characteristics: they have very few unique values relative to the number of samples and the ratio of the frequency of the most common value to the frequency of the second most common value is large. nearZeroVar has been applied to the extracted features using its default parameters. Interestingly, less than 2% of the extracted features (2343 out of 175062) have been selected as informative ones to construct the prediction models (see the supplementary file for more details).

### 3.2 Prediction performance

The performance results of the 227 classifiers with accuracy >50% in the independent test set have been shown in the supplementary figures S1-S3. We have selected the three best classifiers to be included in the AntAngioCOOL package (figure 2): the most sensitive classifier (rpartCost with 88% sensitivity), the most accurate classifier (PART with 75% accuracy) and the classifier with the highest specificity (DeepBoost with 77% specificity). Availability of these three classifiers can help biologists with different questions in mind; e.g. having a list of candidate peptides, what is the narrow list of confident anti-angiogenic peptides or what is the more extended sub-list of candidate anti-angiogenic peptides that contains almost real anti-angiogenic peptides.

**Figure 2:**
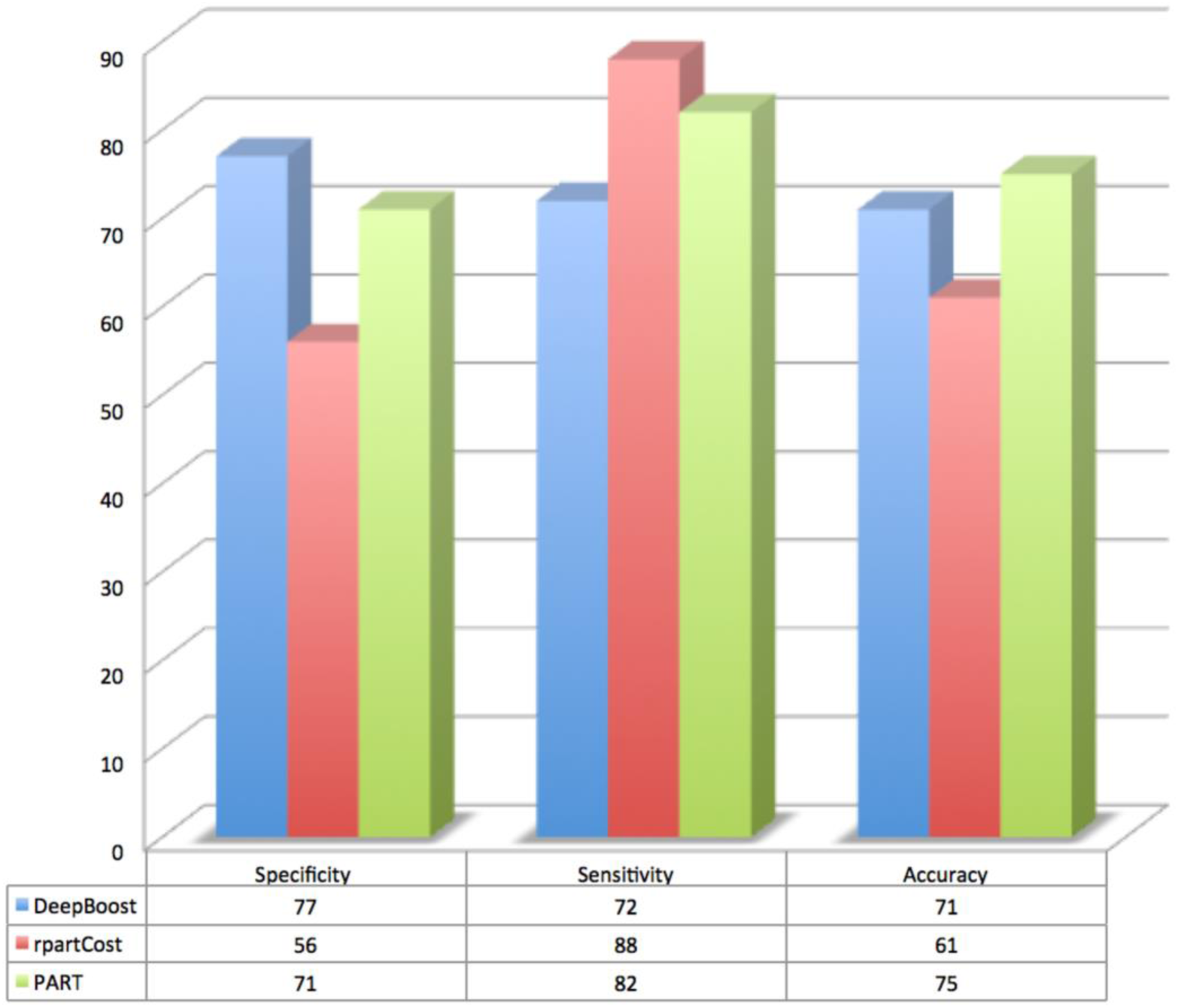
Prediction performance of the three selected classifiers among 227 classifiers to be included in AntAngioCOOL package in the test dataset.

### 3.3 Feature importance

#### physico-chemical profile is the most important feature type

Physico-chemical profile is the main feature type in the final selected set of features: 82% of the final features (figure 3.A). Interestingly, almost all physico-chemical profile features have been selected (1909 out of 1910). Dipeptide and tripeptide compositions are the other important feature types that comprise 9% (200 features) and 4% (101 features) of the final features, respectively. Moreover, figure 3.B shows the percentage of each feature type that has been selected as a subset of the final features.

**Figure 3:**
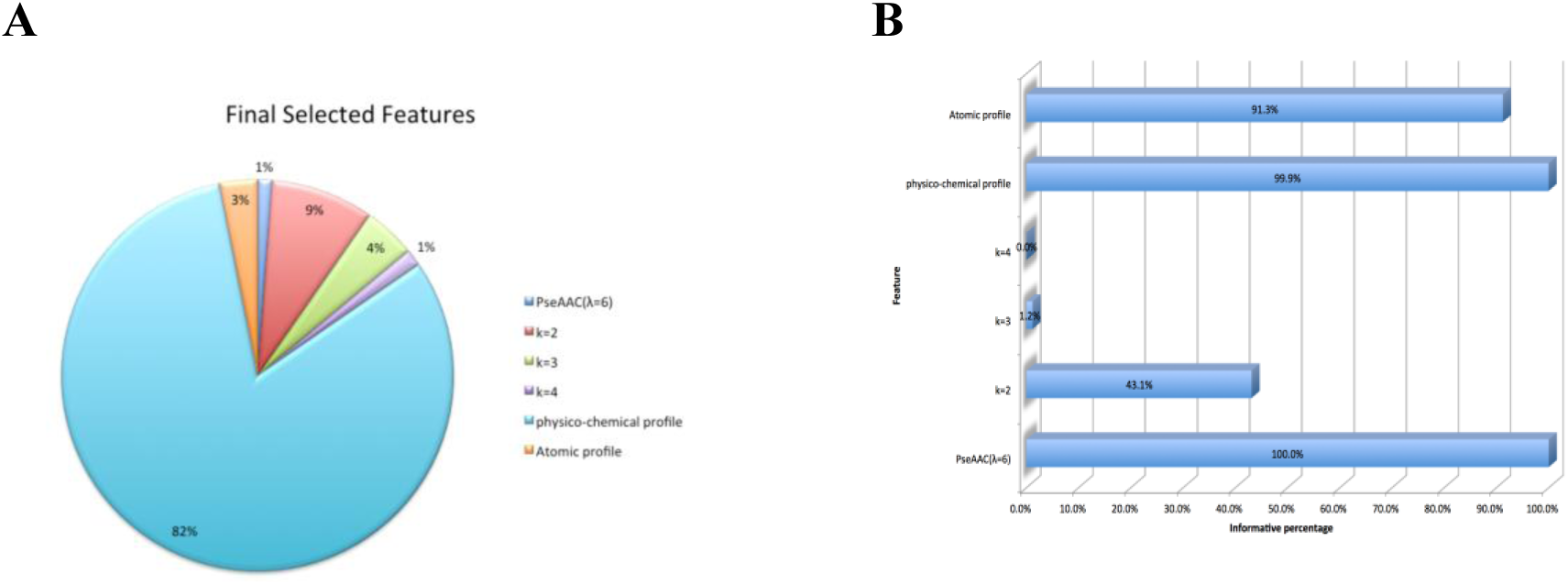
Feature importance according to their contribution in final selected features (informative features). A. Pie chart of the distribution of final features in different feature types. B. The percentage of each feature type that has been selected as informative features.

#### sequence-order information is important in anti-angiogenic peptide prediction

As the figure 3.B shows, in addition to the physico-chemical profile, a considerable percentage of the atomic profile (91.3%) and all the PseAAC features have been selected as informative features (see the supplementary file for more details). One of the important common aspects of these three feature types is that they consider the sequence-order information of the peptide into account. Therefore, this results stress out that the sequence-order information is an effective factor in anti-angiogenic peptide prediction.

#### dipeptide is the most important feature among k-mer composition features

One of the interesting obtained results is that 43.1% of dipeptides have been selected as informative features while for tripeptides and quadpeptides there are very small number of informative features for predicting anti-angiogenic peptides: 101 out of 8512 (1.2%) and 32 out of 164096 (0.02%), respectively. So, dipeptide composition is the most important k-mer composition in anti-angiogenic peptide prediction.

### 3.4 omparison with the current state-of-the-art methods

The proposed method has been trained and tested with the same data used for AntiAngioPred 8. Results reveal that AntAngioCOOL has a higher accuracy (77% vs. 75%) and considerable higher sensitivity (88% vs. 65%). Therefore, AntAngioCOOL package can be used more effectively in anti-angiogenic peptide prediction, especially when one is interested in detecting almost anti-angiogenic peptides (in the cost of having some false positives) in a given list of peptides.

## CONCLUSION

In this study an R package (AntAngioCOOL) has been proposed to predict anti-angiogenic peptides. AntAngioCOOL exploits five descriptor types for a peptide of interest to perform the prediction including: PseAAC, k-mer composition, k-mer composition (reduced alphabet), physico-chemical profile and atomic profile. After removing the non-informative descriptors, only 2% of the extracted descriptors have been used to build the predictor models. AntAngioCOOL includes three different models that can be selected by the user.

The results disclosed that physico-chemical profile is the most important feature type. Also, atomic profile and PseAAC are the other important features. Therefore, it can be concluded that sequence-order information plays a critical role in anti-angiogenic peptide prediction. In addition, according to the results dipeptide has the most contribution in anti-angiogenic peptide prediction among the k-mer composition features.

## Research involving Human Participants and/or Animals

This article does not contain any studies with human participants performed by any of the authors.

